# Chemotactic synthetic vesicles: design and applications in blood brain barrier crossing

**DOI:** 10.1101/061325

**Authors:** Adrian Joseph, Claudia Contini, Denis Cecchin, Sophie Nyberg, Lorena Ruiz-Perez, Jens Gaitzsch, Gavin Fullstone, Xiaohe Tian, Juzaili Azizi, Jane Preston, Giorgio Volpe, Giuseppe Battaglia

**Author notes:** These authors have contributed equally (Adrian Joseph, Claudia Contini, Denis Cecchin). Corresponding author: Prof Giuseppe Battaglia, Christopher Ingold Building, University College London, 20 Gordon 12 Street, WC1H 0AJ, London, United Kingdom.

## Abstract

In recent years, scientists have created artificial microscopic and nanoscopic self-propelling particles, often referred to as nano- or micro-swimmers, capable of mimicking biological locomotion and taxis. This active diffusion enables the engineering of complex operations that so far have not been possible at the micro- and nanoscale. One of the most promising task is the ability to engineer nanocarriers that can autonomously navigate within tissues and organs, accessing nearly every site of the human body guided by endogenous chemical gradients. Here we report a fully synthetic, organic, nanoscopic system that exhibits attractive chemotaxis driven by enzymatic conversion of glucose. We achieve this by encapsulating glucose oxidase — alone or in combination with catalase — into nanoscopic and biocompatible asymmetric polymer vesicles (known as polymersomes). We show that these vesicles self-propel in response to an external gradient of glucose by inducing a slip velocity on their surface, which makes them move in an extremely sensitive way towards higher concentration regions. We finally demonstrate that the chemotactic behaviour of these nanoswimmers enables a four-fold increase in penetration to the brain compared to non-chemotactic systems.

## Introduction

Directional locomotion or taxis is possibly one of the most important evolutionary milestones, as it has enabled many living organisms to outperform their non-motile competitors. In particular, chemotaxis (i.e. the movement of organisms either toward or away from specific chemicals) [6, 7] is possibly the most common strategy adopted by many unicellular organisms to gather nutrients, escape toxins [8] and help coordinate collective behaviours such as the formation of colonies and biofilms [9]. Chemotaxis is also exploited by multicellular systems for tissue development [10], immune responses [11] or cancer metastasis [12]. It enables long-range interactions that extend over length scales that are several orders of magnitude larger than the motile system itself [13]. It is not surprising that scientists have been trying to design devices that mimic such a behaviour [1, 2, 3, 4]. When swimming is scaled down to the microscale, the fluid dynamics are dominated by viscous rather than inertial forces (i.e. Stokes regime). In such conditions, propulsion is possible only by not-time-reversible deformations of the swimmer’s body [14, 15] or by inducing a phoretic slip velocity on the swimmer’s surface [16, 17]. The latter can, for example, be achieved by creating thermal gradients (thermophoresis) or chemical gradients of either charged (electrophoresis) or neutral (diffusiophoresis) solutes in the swimmer’s environment [16]. Recently it has in fact been proposed that the swimmer can induce a slip velocity on its surface by generating an asymmetric distribution of reaction products that creates a localised chemical gradient. This concept known as self-diffusiophoresis was formalised theoretically [18] and demonstrated experimentally using latex particles [19] and gold/silver rods [20].

From a biotechnological point of view, self-propulsion can be applied to create carriers able to autonomously navigate within biological fluids and environments. This could enable directed access to nearly every site of the human body through blood vessels, independent of the blood flow and local tissue architectures. To this respect, recent preliminary experiments were performed with inorganic micro-particles propelled by pH in the stomach of living mice [21]. The ability to control active diffusion as a function of a physiological stimulus bodes well for tackling challenges in drug delivery where an efficient approach is yet to be found. Among these, the ability to deliver drugs within the central nervous systems (CNS) is one of the most difficult tasks where current approaches only enable small percentage of the injected dose to reach the brain and the spinal cord [22, 23]. The brain and the rest of the CNS are well guarded by physiological barriers, with the blood brain barrier (BBB) being the most important. The BBB has the dual function to protect the CNS and to ensure it receives an enhanced supply of metabolites. The brain is indeed the most expensive organ in our body [24] consuming almost 20% of oxygen and glucose. The latter is possibly one of the most important CNS nutrient [25] and the BBB regulates its passage very effectively, with a consequent high flow of glucose from the blood to the brain.

Here we propose the design of an autonomous nanoscopic swimmer based on the combination of naturally occurring enzymes with fully biocompatible carriers, known as polymersomes, that have already proven to hold great promise as drug and gene delivery vehicles [26, 27]. Specifically, in order to target the BBB and enter the CNS [28], we equip polymersomes with the ability to self-propel in the presence of glucose gradients.

## Results and discussion

### Asymmetric polymersomes

Polymersomes are vesicles formed by the self-assembly of amphiphilic copolymers in water [29]. They have been proposed as an alternative to liposomes (vesicles formed by naturally occurring phospholipids) as they offer greater flexibility over chemical and physical properties, and allow large amounts of biological molecules, alone and in combination including proteins and nucleic acids, to be compartmentalised into nanoscale reactors [30, 31]. Furthermore, we have demonstrated [32, 33, 34, 35] that, when two different copolymers are used to form one polymersome, the resulting membrane segregates laterally into patterns whose topology is strictly controlled by the molar ratio of the two copolymers and eventually coarsen into two separate domains forming asymmetric polymersomes [36]. In this article, we exploit this asymmetry to achieve propulsion at the nanoscale. We mixed either poly((2-methacryloyl) ethyl phosphorylcholine)-PDPA (PMPC-PDPA) or poly[oligo(ethylene glycol) methyl methacrylate]-poly(2-(diisopropylamino)ethyl methacrylate) (POEGMA-PDPA) with poly(ethylene oxide)-poly(butylene oxide) (PEO-PBO) copolymers. The copolymers were selected on three different complementary properties: protein resistance for the hydrophilic blocks PEO, POEGMA and PMPC to hinder unspecific interaction with plasma proteins (opsonisation) and limit rapid riddance from the immune system, pH sensitivity for the PDPA to allow endosome-escape and intracellular delivery and finally high permeability for the PBO to preferentially channel both enzyme substrate and product diffusion. PMPC-PDPA and POEGMA-PDPA have been established *in vivo* [37, 38] and while the PMPC can be used directly to target scavenger receptor B overexpressed in cancer cells, [39], the POEGMA is inert in biological fluids and allows easy conjugation to decorate polymersome with ligands [28, 40] to target specific cells. More relevantly here, we showed that we can use POEGMA polymersomes as platform for crossing the BBB and entering the CNS when combined with the low density lipoprotein receptor-related protein 1 (LRP^-1^) targeting peptide Angiopep-2 (LA) [28]. PEO-PBO forms very thin membranes (∼ 2.4 nm) [41] that are highly permeable to most small polar molecules, such as hydrogen peroxide and glucose [42]. The schematics of our proposed design is shown in Fig. 1a. We have previously demonstrated [33] that the two copolymers form asymmetric polymersomes at an optimal 9:1 molar ratio with the small permeable bud being formed by the minor PEO-PBO component. This can be verified using transmission electron microscopy (TEM) by imaging the polymersomes using positive staining selective for the PDPA blocks (see Figs. 1b-c for the PMPC-PDPA/PEO-PBO and the POEGMA-PDPA/PEO-PBO mixtures respectively). As shown using negative staining TEM (Fig. 1d) where the PBO domain is darker, the thickness of the two membranes can be measured to be about 6.4 nm and 2.4 nm confirming previously reported measurements [26, 41].We have already demonstrated that PBO membranes are 10 times more permeable than phospholipid ones [42] and that these are at least 10 times less permeable than thick membranes formed by aliphatic chains such as the PDPA ones [43]. To a first approximation, we can thus infer that the PBO membrane is two orders of magnitude less permeable than the PDPA membrane. We can employ such an asymmetric polymersome to encapsulate enzymes using a technique based on electroporation [31]. We chose glucose oxidase to catalyse the glucose oxidation to form d-glucono-J-lactone and hydrogen peroxide and catalase to catalyse the decomposition of hydrogen peroxide into water and oxygen. Both enzymes and reagents are naturally occurring in the human body. As shown in Supplementary Table 1 and Supplementary Fig. 1, we encapsulated an average of 6 glucose oxidases and 2 catalases per polymersome either alone or in combination. We thus hypothesise that, as the enzymes react with their respective substrates, the confined reactions will produce a flux of products that will be preferentially expelled out of the polymersomes from the most permeable patch, i.e. the bud formed by the minor PEO-PBO component. This in turn generates a localised gradient of the products that should set up the conditions for self-propulsion. The nature of the propulsion mechanism depends on the interaction between the reaction products and the two different polymersome domains [16]. To a first approximation, this should set the conditions for*self-diffusiophoresis* where the depletion of the product molecules near the polymersome surface induces a lateral water flow with slip velocity, **v_S_**. Assuming a spherical geometry of radius *R*, the polymersome propulsion translation and angular velocity can be derived form the slip velocity as 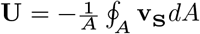 and 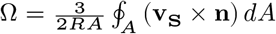 respectively, with *A* being the total polymersome surface area and **n** the polymersome orientation unit vector. This vector originates from the polymersome centre of mass and its directed toward the centre of the asymmetric PEO-PBO domain. Both velocities can be used to derive the general equations of motion expressed as a function of the polymersome position **r** and orientation unit vector **n** as:

**Figure 1:**
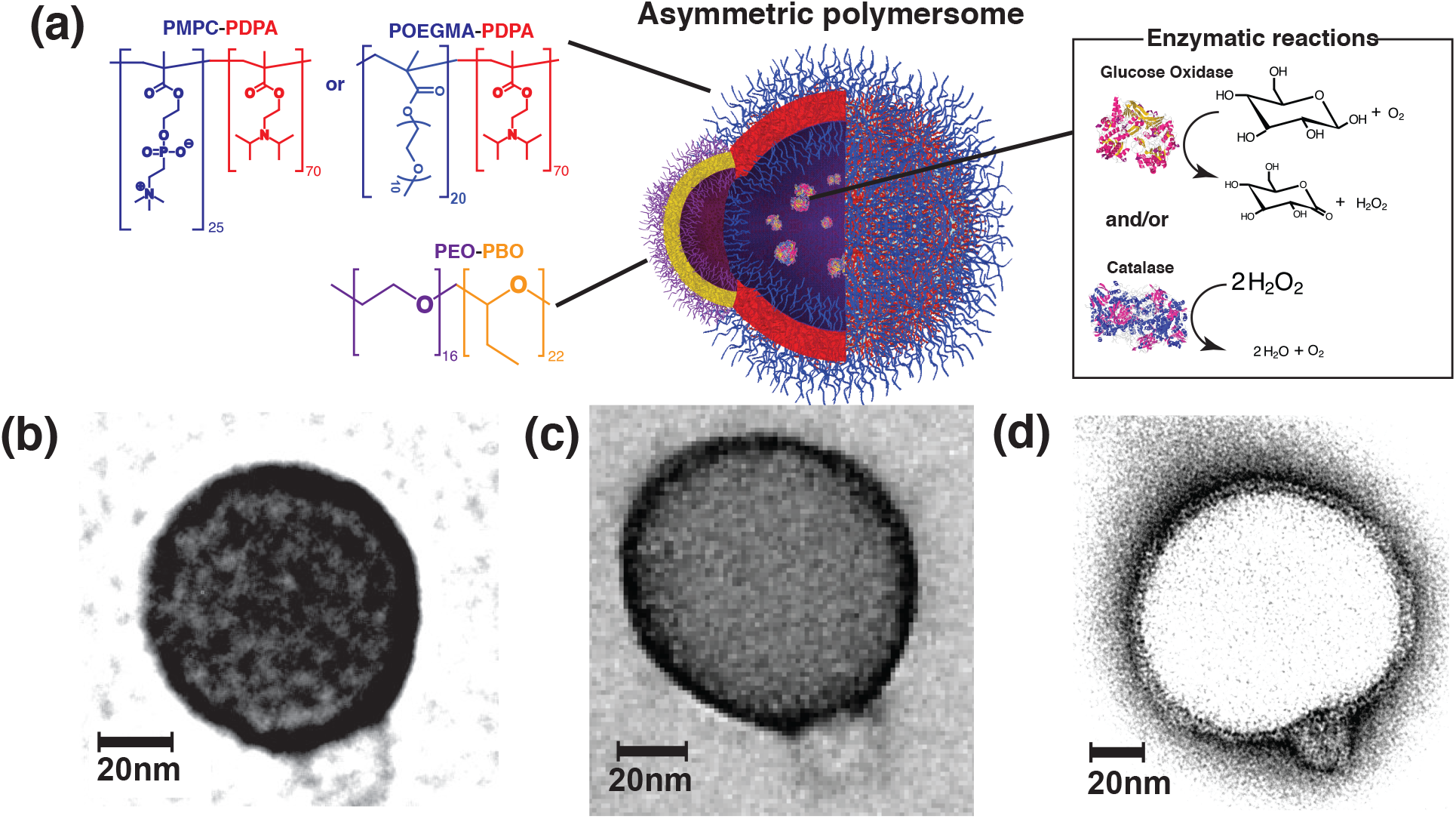
Asymmetric polymersomes. (a) Schematic representation of a chemotactic polymersome using a combination of membrane topology formed by PEO-PBO copolymers mixed either with PMPC-PDPA or POEGMA-PDPA copolymers. The polymersomes encapsulate glucose oxidase and/or catalase enzymes.**(b)** 9:1 PMPC-PDPA/PEO-PBO polymersome imaged in positive staining exploiting the high affinity of PDPA with the staining agent phosphotungstic acid (PTA).**(c)** 9:1 POEGMA-PDPA/PEO-PBO polymersome imaged in the same staining agent for PDPA.**(d)** 9:1 PMPC-PDPA/PEO-PBO polymersome imaged in negative staining to highlight the differences in membrane thickness between the PDPA and the PBO membrane.

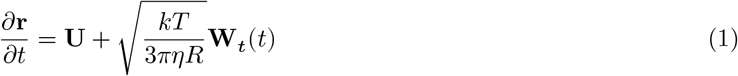

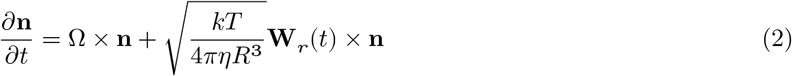

where *k* is the Boltzmann’s constant, *T* the absolute temperature, η the water viscosity, **W**_*t*_ and **W**_*r*_ are white noise vectors that respectively model the translational and rotational Brownian diffusion of the particle [5, 16].

### Active diffusion analysis

To characterise the motility of the polymersomes, we have employed a technique known as nanoparticle tracking analysis (NTA) [44]. This is based on the dark-field parallel tracking of thousands of single nanoparticles using a camera to detect the light of a monochromatic laser scattered by the particles. The geometry of the observation chamber is shown in Supplementary Fig. 2 and, unless specified differently, we performed all the measurements atphysiological conditions, i.e. T = 37 °C, ɳ_water_ = 0.69 mPas, pH=7.4, in 100 mM phosphate buffer solution (PBS). The trajectories and the corresponding mean square displacements (MSDs) can be used to evaluate the motility of the polymersomes. In Supplementary Figs. 3–13 we show 1-s trajectories (all normalised to a common origin) and the corresponding MSDs for thousands of polymersomes imaged at 30 frames per second (fps) under different environmental conditions. In a homogeneous environment, either in presence or absence of the substrate, the results show that, independently of being symmetric or asymmetric, loaded with enzymes or empty, the polymersomes have a typical Fickian diffusion profile with linear MSDs and stochastic trajectories. While the MSDs averaged over thousands of trajectories (Supplementary Figs. 3–7) show some variations in the long-time diffusion coefficient, these variations are mainly due to statistical fluctuations between different experimental realisations of the process. In particular, we do not observe any appreciable enhancement in diffusivity. This suggests that even if the enzymatic reaction creates an asymmetric distribution of products around the loaded patchy polymersomes, with consequent propulsion velocity, any corresponding directed part of the motion is not sufficient to overcome the polymersome high rotational diffusion due to its small size (z-average measured by DLS *R =* 50 ± 10 nm, Supplementary Fig. 1), which effectively hinders any self-propulsion by effectively randomising the particles’ orientations in τ ≈ 0.5 ms, one order of magnitude below our experimental time resolution (about 33.3 ms). To further confirm this, we calculated the ratio between the enhanced diffusion coefficient *D_eff_* and the Stokes-Einstein diffusion coefficient*D_0_* from 2D projections of 3D simulated trajectories of chemotactic polymersomes If in first approximation we assume that Ω = 0, for a polymersome moving with a propulsion velocity of 100μ*ms*^-1^, *D_eff_* = 1.15*D*_0_, for a polymersome moving at *200μms*^-1^ the *D*_eff_ = 1.45*D*_0_ A detectable enhancement o*D*_eff_ = 2*D*_0_ corresponds to a propulsion velocity of *300μms^-1^*. These calculations confirm that any enhancement in diffusion is small for realistic values of size and velocity in our system, thus making it difficult to detect given the experimental variability. Both experiments and simulations therefore suggest that the angular phoretic term proportional toΩ in equation 2 is considerable smaller than the Brownian rotational component and hence can be ignored hereafter.

We repeated the same set of experiments of Supplementary Figs. 3–7 in the presence of a concentration gradient created by adding the substrate from one side of the observation chamber (Fig. 2 and Supplementary Figs. 8–16). Under the new experimental conditions, the symmetric polymersomes (either loaded or empty) as well as the empty asymmetric polymersomes still showed a typical Fickian diffusion profile with stochastic trajectories and linear MSDs as a function of time. As a reference, Fig. 2a only shows the data corresponding to the case of symmetric polymersomes loaded with both glucose oxidase and catalase responding to a gradient of glucose (generated by a 1M-solution at the injection site), while the other control measurements are reported in Supplementary Figs. 3–16. The enzyme-loaded asymmetric polymersomes instead responded quite differently to the gradient of their respective substrate (Figs. 2b-e). Fig. 2b shows the data for the asymmetric polymersomes loaded with catalase alone (Cat) responding to a hydrogen peroxide gradient (generated by a 1mM-solution) coming from the right-hand side of the observation chamber; the normalised trajectories are biased toward the gradient and the corresponding MSDs show a ballistic behaviour with a quadratic dependence on time. We limited our experiments to low concentration of hydrogen peroxide to avoid its spontaneous decomposition and consequent formation of oxygen bubbles that could dissolve the polymersomes. Such a super-diffusive behaviour is considerably more pronounced for the asymmetric polymersomes loaded either with glucose oxidase alone (Gox) (Fig. 2c) or glucose oxidase and catalase (Gox+Cat) together (Fig. 2d-e) responding to a glucose gradient generated by a 1M solution; almost all the trajectories are aligned toward the gradient, whether this comes from the right-(Fig. 2d) or the left-hand side (Fig. 2e). This does not change when instead of using PMPC-PDPA polymersomes we use POEGMA-PDPA polymersomes demonstrating that the differential permeability of PDPA and PBO are responsible for the self-phoresis (Fig. 2f). In addition to the trajectory and MSD analysis, the average drift velocities are plotted in Fig. 2g as a function of the time of observation after the substrate addition. For Brownian particles such as those in the control samples, the average drift velocity is zero but, as the samples become more chemotactic, the drift velocity gradually increases. The variation of the drift velocity as a function of time after the addition of the substrate allows us to estimate how the self-propulsion behaviour varies with the chemical gradient magnitude, and, in all cases the drift velocity equilibrates to a plateau value corresponding to the time when the gradient becomes linear (i.e. ∇*C* ≈ constant) and the system reaches steady-state conditions. Finally, the distribution of the particle orientation with respect to the direction of the substrate gradient is plotted in Fig. 2h for all cases. Brownian samples (such as all the controls) have directions almost equally distributed across all angles, while, as the sample starts to exhibit propulsion and chemotaxis, the distribution of particles polarises toward the direction of the gradient. All the data displayed in Fig. 2 show that independently of the enzyme/substrate system, asymmetric polymersomes show typical ballistic behaviour with a chemotactic response towards the enzyme substrate gradient marked here asθ = 0. The catalase-loaded polymersomes respond rather weakly to the hydrogen peroxide gradient and this is independent of the peroxide initial tested concentrations. Glucose oxidase-loaded asymmetric polymersomes, on the other hand, respond very strongly to a glucose gradient, reaching drift velocities around 20 μm s^-1^ with most particles polarised toward the gradient. Interestingly, similar values are comparable to those of chemotactic bacteria, such as *E. coli*, which are one order of magnitude larger than the polymersomes studied herein [9]. As shown in Fig. 1a, glucose oxidase and catalase operate very well together as their respective reactions feed each other with hydrogen peroxide being a product of glucose dissociation and the oxygen being a product of hydrogen peroxide dissociation [45]. Furthermore, their combination leads to the formation of non-detrimental molecules as both oxygen and hydrogen peroxide are consumed and transformed into water and d-glucono-δ-lactone. Glucose oxidase self-regulates and, as a critical concentration of hydrogen peroxide is reached, its activity is inhibited. This means that even at low H_2_O_2_ concentrations, we can assume that catalase consumes most H_2_O_2_ [46]. Most notably, glucose oxidase and catalase loaded asymmetric polymersomes had the strongest response to glucose gradients, and indeed produced slightly higher drift velocities and considerably more polarised chemotaxis than the system loaded with glucose oxidase alone or catalase alone. From these data we can conclude that no osmotic flow is generated as demonstrated by all control measurements in the supplementary figures and that: *(i)* the asymmetric distribution is critical, indeed symmetric polymersomes (either made of PDPA or PBO membranes) loaded with enzymes did not show any chemotactic drift;*(ii)* the reaction is critical, and empty polymersomes either symmetric or asymmetric do not exhibit any diffusophoretic drift due only to the substrate gradient; and finally *(iii)* only when the enzymes are encapsulated within an asymmetric polymersome chemotaxis is exhibited suggesting that the propulsion velocity is only proportional to the products ∇*C_p_*. These conclusions suggest that, in the presence of a gradient, the strength of the polymersomes’ propulsion velocity is strongly biased by its orientation so to create an asymmetric angular probability in the particle’s motion that is higher when the particle is oriented toward the gradient. The data in Fig. 2 are the 2D projections of 3D trajectories on the field of view plane. In order to simulate the same arrangement, we use a spherical polymersome with *R* = 50nm and a smaller semi-spherical patch radius, *r* = 15nm as shown in Fig. 3a. We assume that the chemical gradient is aligned along the **x**-axis and that the orientation of the unit vector, is defined by a cone within the sphere with aperture*2β*. We can simulate the distribution of the products’ concentration just outside the PBO permeable patch at different orientations β (Supplementary Note 3.4.2) and this is expectedly biased toward the chemical gradient as shown by the red line in Fig. 3a. We approximated this distribution with the function 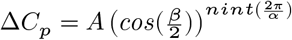, where *A* is a proportionality constant and *α* is the sector angle of the PBO domain. Since the gradient in the product distribution around the particle is in first approximation proportional to such ∆*C_p_* [18, 19], the propulsion velocity can be estimated from the data by describing its functional form with the same modulation in the polymersome orientation. In fact, such an approximation, together with the assumption that the polymersome’s phoretic angular velocity Ω is negligible when compared to its rotational diffusion, allow us to simulate the propulsion of the polymersomes in the presence of the substrate gradient by using equations 1 and 2 (Supplementary Note 3.3). As shown in Fig. 3b, by fitting (solid lines) the experimental data (circles) with our model, we were able to estimate the strength of the propulsion velocity for each formulation (Fig. 3c). The (Gox + Cat) formulation is the one with the highest propulsion velocity and the formulation with catalase alone in the presence of hydrogen peroxide the one with the lowest value. The difference in performance of the two enzymes/substrates is possibly due to the difference in substrate concentration (considerably lower for the peroxide)which lead to a shallower gradient of products. Notably, we observe chemotaxis in all the different combinations proving that the system we propose here works with very different combinations of substrate/enzyme. More importantly, the simulations allow us to access the dynamics of propulsion with no limits in both spatial and temporal resolution. In Fig. 3d we show the simulated 3D trajectories normalised to a common origin of 20 polymersomes with a temporal sampling identical to our experimental setting (i.e. 33ms corresponding to a 30 fps acquisition rate). We show these both as 3D axonometric projection and in the corresponding*xy* plane view which reproduce very closely the experimental data in Fig. 2. In Fig. 3e, a single trajectory is plotted using both temporal resolution of 33ms (blue line) and 33μs (orange line) corresponding to a 30 and 3.10^5^ fps acquisition rate respectively. The polymersome trajectories reveal that they are the result of a succession of running and re-orientation events within the ms timescale and hence the polymersomes quickly re-orient toward the gradient with consequent self-propulsion as schematically represented in Fig. 3f.

**Figure 2:**
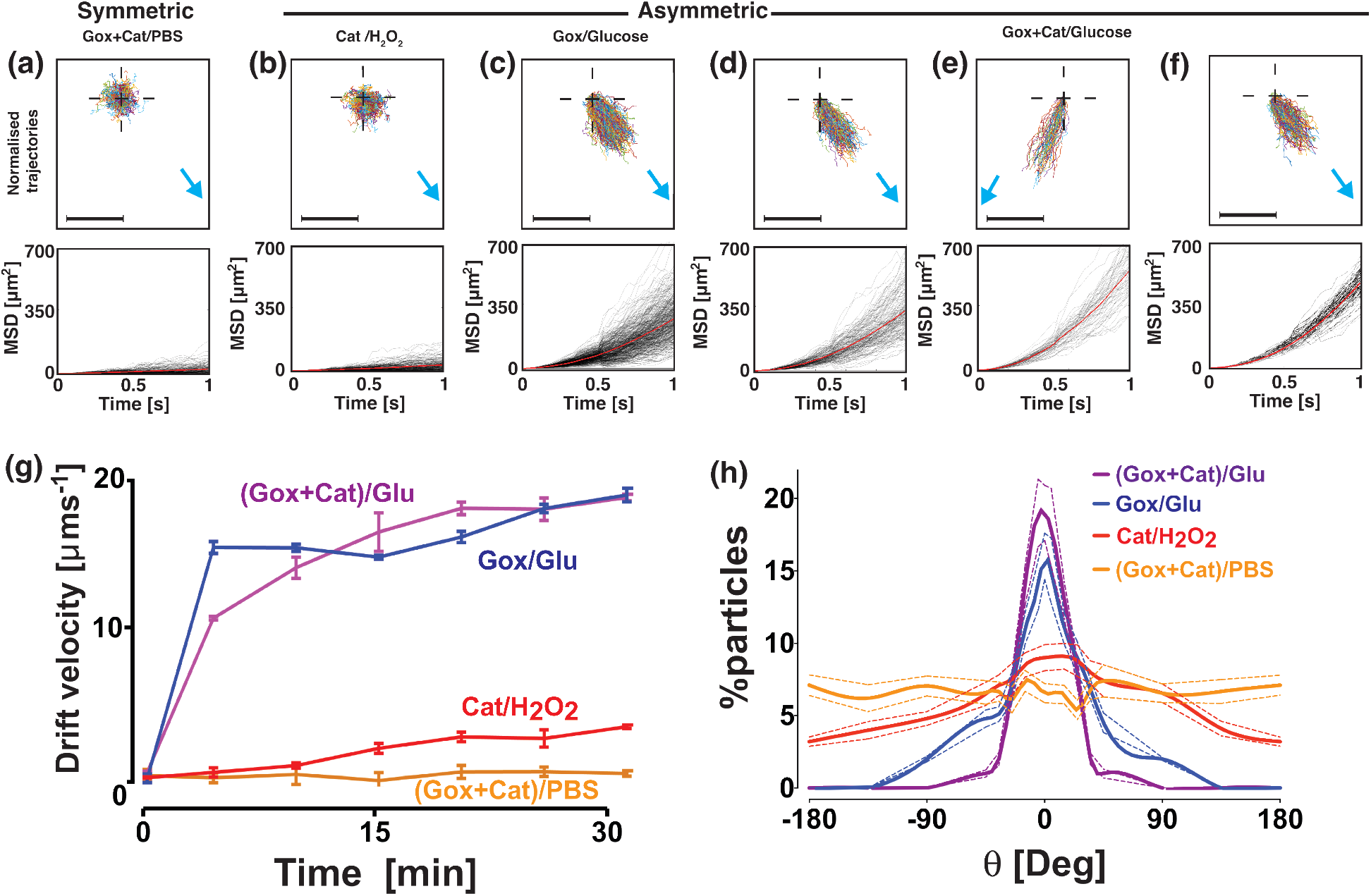
Single particle analysis in the presence of a chemical gradient. Normalised 1s-trajectories (the cross marks the common origin) and corresponding mean square displacements (MSDs) for **(a)** symmetric PMPC-PDPA polymersomes loaded with glucose oxidase (Gox) and catalase (Cat) and responding to a glucose gradient, **(b)** asymmetric PMPC-PDPA/PEO-PBO polymersomes loaded with catalase and responding to a hydrogen peroxide gradient, **(c)** loaded with glucose oxidase and responding to a glucose gradient, **(d-e)** loaded with glucose oxidase and catalase responding to a glucose gradient coming **(d)** from the right-hand side and **(e)** from the left-hand side and for **(f)** asymmetric POEGMA-PDPA/PEO-PBO polymersomes loaded with glucose oxidase and catalase responding to a glucose gradient coming from the right-hand side. The scalebar is 20 μm, and the blue arrows indicate the direction of the substrate gradient. **(g)** The average drift velocity is plotted as a function of time after the substrate addition for the previous experiments.The error bars represents the standard error calculated over n=3 measurements **(h)** Degree of polarisation of the corresponding trajectories towards the chemical gradient plotted as percentage of particles versus the gradient angle. Perfect alignment with the gradient corresponds to θ = 0 degrees. The dashed lines represent the standard errors.

**Figure 3:**
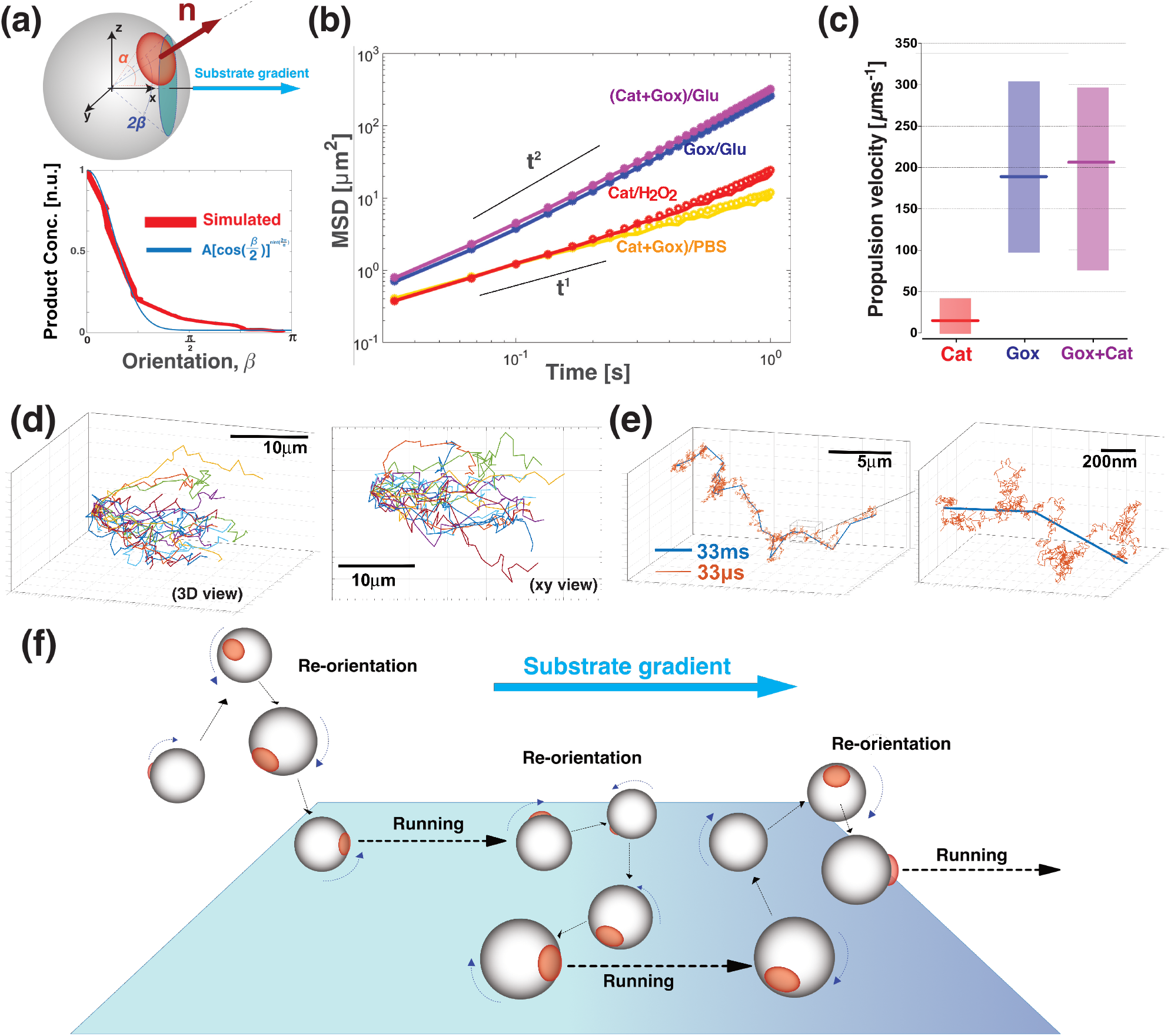
Polymersome chemotaxis simulations. (a) Schematics of an asymmetric polymersome and its reference axis. **W**e assumed the polymersome to be a sphere (*R* = 50 nm) with a smaller patch (*r* = 15 nm and sector angle α); the angle β represents the orientation of the unit vector **n** with respect to the chemical gradient **∇C** here aligned to the *x*-axis. **W**e simulated the distribution of the products around the polymersome and their normalised concentration is plotted (red line) alongside a fitting function 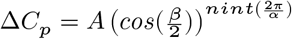 (blue line). **(b)** Average MSDs for both experimental (circles) and simulated data (solid line) for asymmetric polymersomes loaded with Gox and Cat responding to a glucose gradient (purple line and data) or in PBS (orange line and data), loaded with Gox and responding to a glucose gradient (blue line and data) and loaded with Cat responding to a hydrogen peroxide gradient (red line and data). **(c)** Corresponding propulsion velocities calculated by the numerical fittings for the three different combinations of enzymes and substrates.The lines represent the average values while the bars represent the range of minimum and maximum calculated velocity in the sample. **(d)** 20 simulated trajectories of Gox+Cat loaded polymersomes using the same temporal steps as in the experiments (30fps). These are shown as a 3D axonometric projection view and in the corresponding *xy* plane to show the comparison with the experimental data. **(e)** A single simulated 3D trajectory shown with temporal steps of 33ms (blue line) and 33μs (orange line). The detail of a single trajectory is zoomed to show the succession of re-orientation and running steps of the polymersome diffusion. **(f)** Schematics of the proposed mechanisms of asymmetric polymersome chemotaxis, which consists of an alternation of running and re-orientation events.

### Chemotaxis in complex environments

In order to get further insight into the chemotactic response of our system, we performed further experiments on the polymersomes loaded with both enzymes to assess their chemotactic capability more quantitatively using the approach shown in Fig. 4a. A cylindrical agarose gel, pre-soaked in a 1-M glucose solution, was placed on the edge of a Petri dish filled with PBS. Various polymersome formulations were added at the centre of the dish with a syringe pump. Samples were collected at different locations within the Petri dish and at different time points as shown in Fig. 4b, and quantified for concentration and sizing (Supplementary Note 3.2.1 and Supplementary Figure 17). In Fig. 4c-e we show concentration maps of the polymersomes in the dish at time 0 (Fig. 4c) and 10 min after their addition, both for the symmetric formulation (Fig. 4d) and for the asymmetric formulation (Fig. 4e) loaded with glucose oxidase and catalase, in response to a glucose gradient. We also studied a different configuration (Supplementary Note 3.2.2): a Petri dish pre-filled with fluorescent polymersomes where a drop of 1M-glucose solution is added in the centre of the dish, which is directly imaged with a fluorescence camera (Fig. 4f). The corresponding fluorescence images of both symmetric and asymmetric polymersomes before glucose addition and at times t = 0, 10 and 15 min are shown. While the first experiment shows that the asymmetric polymersomes do not dilute in the presence of the glucose gradient, and instead almost entirely drift towards the glucose source (Fig. 4e), in the second experiment we can observe that the asymmetric polymersomes can concentrate towards the glucose gradient from high dilutions (Fig. 4g).

**Figure 4:**
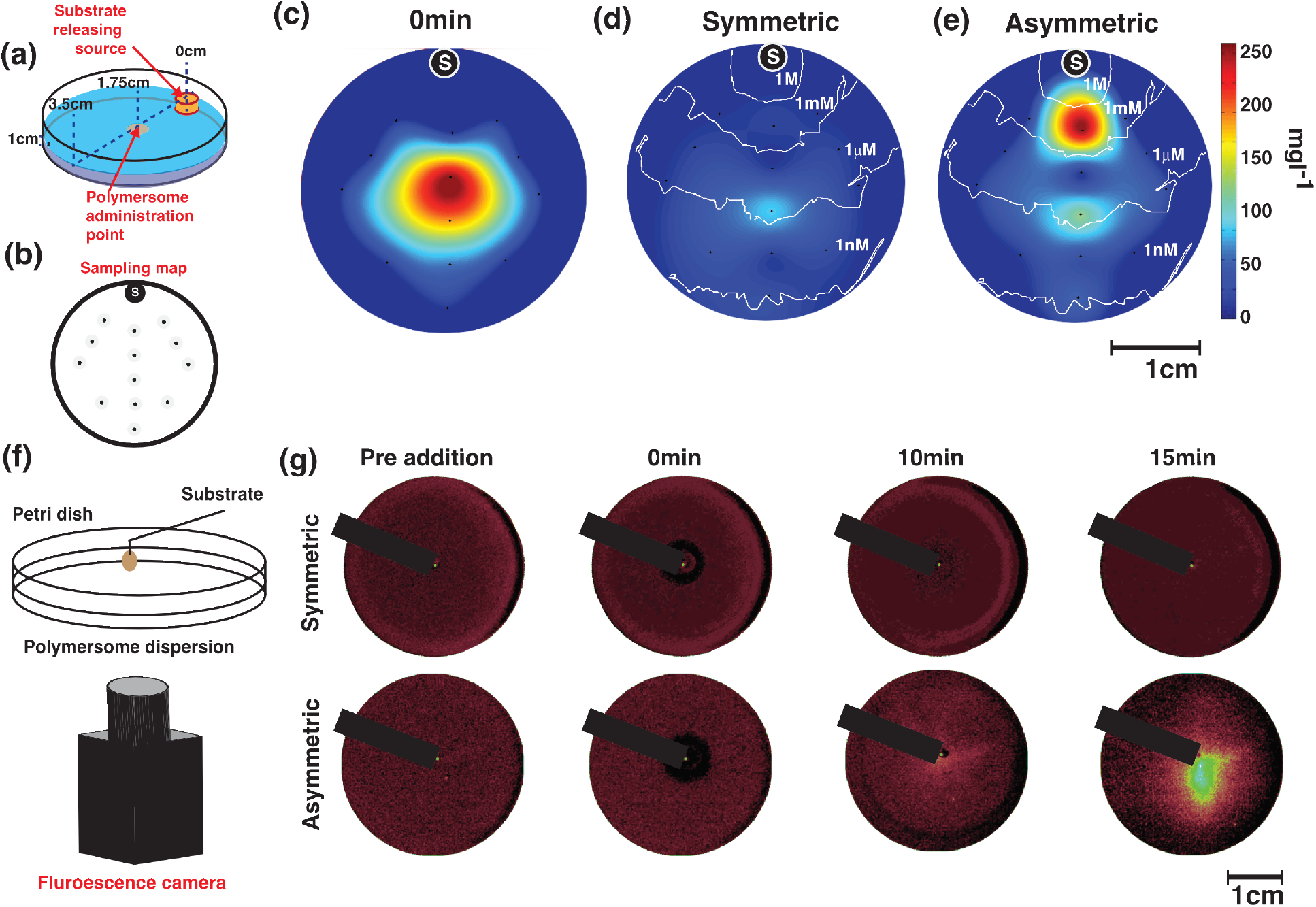
Long-range chemotaxis. (a) Schematics of a Petri dish where a cylindrical agarose gel soaked in glucose is placed. A time t = 0, a 1mg ml^-1^ concentration of polymersomes is added in the dish centre and their concentration is sampled at different locations as indicated by the sampling map in **(b).** The dot labelled with S indicates the position of the source of glucose. **(c-e)** The resulting maps show the two-dimensional distribution of asymmetric polymersomes **(c)** at time t = 0, and the distribution of polymersomes at time t = 10 min for **(d)** symmetrical PMPC-PDPA and **(e)** asymmetrical PMPC-PDPA/PEO-PBO polymersomes loaded with catalase and glucose oxidase. The isocratic white lines show the glucose gradient calculated by computational fluid dynamics. **(f**) A similar experiment is performed by adding glucose in the centre of a Petri dish containing fluorescently labelled polymersomes after they have thermalised in it. The imaging is performed with a fluorescence camera. **(g)** The corresponding fluorescence images are shown for both symmetric PMPC-PDPA and asymmetric PMPC-PDPA/PEO-PBO polymersomes loaded with catalase and glucose oxidase at different times: before the addition of glucose, at time t = 0, 10 and 15 min. The black line indicates the needle for the injection of glucose over the imaging camera.

These experiments show quite convincingly that the chemotactic polymersomes follow shallow gradients and concentrate toward a given chemical source over time scales of minutes and length scales 107 times longer than the swimmer’s characteristic size.

All the data bode well for bestowing polymersomes with chemotactic capability and indeed augmenting their efficiency in navigating across biological barriers. To understand the effect of flow, we performed the same experiments as in Fig. 2 but in the presence of a constant flow almost perpendicular to the glucose gradient. The two chosen flow rates of 0.5 and 3.5 μlmin^-1^, corresponding to velocities of 10 and 150 μms^-1^ (i.e. Péclet number of 0.15 and 2.3 respectively) represent conditions encountered next to the capillary barriers or right in the capillary centre respectively. As shown in Fig. 5a, the normalised trajectories for both pre-substrate addition and symmetric polymersomes show a typical Gaussian distribution that is more skewed as the flow rate increases from 0 to 3.5 μlmin^-1^. At zero flow, the glucose oxidase and catalase loaded polymersomes show a rapid response to the glucose gradient with overall drift plateauing at about 20 min after the addition of glucose. At a flow rate of 0.5 μlmin^-1^, the chemotactic drift is still sufficient large to overcome the convection and indeed polymersomes still move toward the glucose gradient, albeit at lower velocities. At a flow rate of 3.5 μlmin^-1^, the chemotactic drift combines with the flow inducing a drift of the polymersomes with trajectories taking a direction of about 45° from the flow line. It is important to note (as shown in Fig. 5a) that as the flow increases the gradient vector rotates from its original unbiased position to being almost perpendicular to the flow. In order to test the effect of placing chemotactic polymersomes in blood flow, we employed an agent-based model of the nanoparticles in capillaries in the presence of erythrocytes (also known as red blood cells) that we have developed previously [47]. In Fig. 5b, we show a snapshot of the streamlines of the flow observed in a capillary with a radius of 4 μm and length of 800 μm calculated by computational fluid dynamics (Supplementary Note 3.1). The red cylinders represent erythrocytes (at physiological haematocrit H% =10.7%) and the colour maps show the normal velocity, i.e. the velocity component perpendicular to the vessel wall. We used this geometry and we seeded 100 nanoparticles randomly at the entrance of the vessel and allowed their passage through the vessel. The vessel walls were set as no-slip, sticky boundaries (i.e. as a polymersome approaches the barrier it binds to it), so that the number of nanoparticles bound to the vessel wall could be evaluated with different sized particles and velocities of propulsion. As discussed above, we can assume that as asymmetric polymersomes encounter a glucose gradient they will propel with a propulsion velocity that is directly proportional to the gradient, and their rotation is uniquely controlled by Brownian dynamics. Assuming a glucose gradient across the vessel, we performed the calculations for polymersomes with radius *R* = 50, 100, and 250 nm, which is representative of a typical size distribution of polymersomes (see DLS measured distributions in Supplementary Fig. 1), and to represent the spread of propulsion velocities (see both Figs. 2 and 3) we propelled the polymersomes at from 0 to 200 μms^-1^. Fig. 5c shows the percentage of particles that bind to the vessel wall during a single passage. Binding to the vessel walls is generally improved by increasing the propulsion velocity. Indeed propulsion augments binding 2-fold from 0 to 200 μms^-1^ for small nanoparticles and the binding to the wall is considerably improved for the case of larger polymersomes and high propulsion velocity reaching almost 100% of particles binding. Bigger particles bind better to the wall than smaller particles due to their smaller rotational diffusion which keeps the particles’ orientation along the gradient for longer [4]. Modelling would thus suggest that adding an element of propulsion to the motion of the polymersomes increases the overall uptake from the blood due to their improved distribution to the endothelial wall interface. Furthermore, the use of glucose as a substrate ensures that there is a high level of substrate available within the blood, as blood glucose is maintained at 4–7.8 mM [25]. In addition, brain metabolism requires high levels of glucose and glucose transporters are well known to be over-expressed on the BBB [25] and hence it is not far-fetched to assume that blood glucose has a positive gradient toward the blood wall and an even more favourable distribution within the brain. Recently, we have demonstrated that polymersomes can be conjugated with peptides that target the LRP^-1^ receptor. This receptor is over-expressed at the BBB and it is associated with a transport mechanism known as transcytosis. We have demonstrated that by targeting this pathway we can deliver large macromolecules to CNS resident cells [28]. LA modified asymmetric polymersomes can cross the BBB and we showed this using a 3D *in vitro* BBB model that comprises two cell types: brain endothelial cells and pericytes cultured in the presence of conditioned media from astrocytes. The endothelial cells are placed on the upper compartment and they are separated from the pericytes by a porous polycarbonate membrane (pores < 0.4/*mwm*)[28]. The geometry of the model is shown in the Supplementary Fig. 18a alongside with the qualitative (Supplementary Fig. 18b) and quantitative (Supplementary Fig. 18c) kinetics of the polymersomes BBB crossing. These data show effective crossing and active pumping of the LA-polymersomes from the apical to the basolateral side of the BBB performed by the endothelial cells. Moreover, the same *in vitro* model can be used to evaluate the early time points, and as shown in Fig. 5d and Supplementary Figure 19, we observed that LRP^-1^ mediated transcytosis is extremely fast taking about 15s from the binding event on the apical side to a full crossing to the basolateral side. We have here used this system to demonstrate that chemotaxis can indeed augments delivery significantly. This effect was validated in the rat CNS through *in situ* brain perfusion and quantification of fluorescently labelled polymersomes in the different parts of the brain by fractionation. Chemotactic polymersomes, responsive to glucose and functionalised with LA, demonstrated about a 4-fold delivery increase into the parenchyma compared to non-chemotactic polymersome controls, including LA-modified asymmetric empty polymersomes, LA-symmetric polymersomes either loaded with Gox+Cat or empty (Fig. 5e). The effective passage across the BBB is further demonstrated by immune-fluorescence histologies of the brain sections whose capillaries are stained using the CD34 marker (green), the cell nuclei are stained with Hoescth (blue) and the polymersomes are labelled with Cy5 (red) as shown in Fig. 5f. The non-active polymersomes were optimised to reach a respectable 5% of the injected dose. However, modifying the polymersomes, by adding an asymmetric patch and by loading them with glucose oxidase and catalase, enabled a staggering delivery of 20% of the injected dose, which to the best of our knowledge has never been reported so far with any other system. The glucose is a required metabolite in the blood and the brain consumes more than the 20% of the assimilated glucose at any given time. It is also established that the brain endothelial cells express extremely high level of glucose transporters [48] suggesting that as the blood reach the brain area, there must be a gradient from the centre to the wall of the vessel.

**Figure 5:**
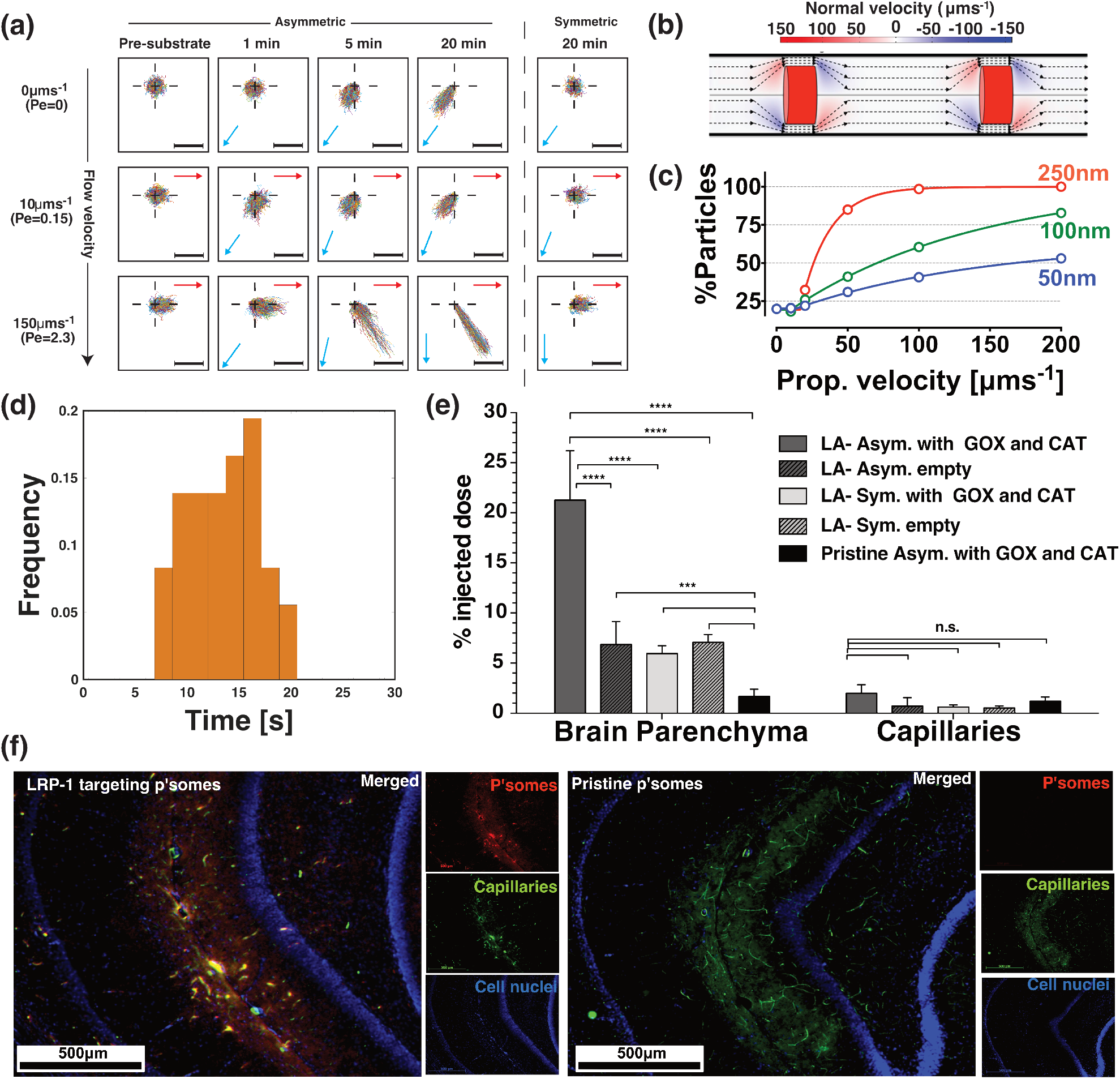
Chemotaxis under flow and *in vivo*. (a) Normalised polymersome 1-s trajectories measured in the presence of steady state flow (0, 0.5 and 3.5 μms^-1^) and collected prior, 1, 5 and 20 min after the glucose gradient addition for both PMPC-PDPA/PEO-PBO asymmetric and PMPC-PDPA symmetric polymersomes loaded with glucose oxidase and catalase. The scalebar is 20 μm, and the red arrows denote the direction of the flow within the observation area while the blue arrows denote the average direction of the glucose gradient within it. **(b)** Streamlines of flow observed in a capillary with radius of 4 μm and length of 800 μm calculated by CFD. The red cylinders represent erythrocytes (haematocrit H% =10.7%) and the colour map shows the normal velocity of the flow, i.e. the component perpendicular to the vessel walls. **(c)** Simulated percentage of the total number of particles bound to the vessel surface as a function of their drift velocity in a gradient for 50, 100 and 250 nm asymmetric nanoparticles calculated with an agent-base model of chemotactic particles within a capillary such as in **(b**)Note: the error bars show the standard error. **(d)** Frequency distribution of the crossing time from apical to basolateral of LA-POEGMA-PDPA polymersomes measured over 35 different measurements using the *in vitro* BBB model as showed in Supplementary Figure 18 (note one example measurement is showed in Supplementary Figure 19). **(e)** Percentage of the injected dose found in the rat brain parenchyma and the capillary fraction 5 min after intra-arterial injection of LA-POEGMA-PDPA/PBO asymmetric polymersomes loaded with Gox+Cat and empty, and LA-POEGMA-PDPA symmetric polymersomes loaded with Gox+Cat and empty, as well as pristine asymmetric POEGMA-PDPA/PEO-PBO polymersomes loaded with Gox and Cat (n=6. Statistical significance: *** *p* < 0.001 and **** *p* < 0.0001). The error bars show the standard error. **(f)** Immunefluorescence histologies of rat hippocampus sections of animals treated with LA-POEGMA-PDPA/PBO asymmetric polymersomes loaded with Gox+Cat and pristine asymmetric POEGMA-PDPA/PEO-PBO polymersomes loaded with Gox and Cat.

## Conclusions

We have shown here that an established intracellular delivery system such as PMPC-PDPA and POEGMA-PDPA polymersomes can be modified to possess chemotactic capabilities toward glucose gradients. We achieve this by using a novel process of converting a chemical potential difference into an actual propulsion mechanism capable of tracking small molecule gradients over distances that are many orders of magnitude greater than the nanoparticle’s characteristic length. We demonstrated that nanoscopic polymersomes move according to superdiffusional behaviours and most importantly they do so only in the presence of a gradient becoming chemotactic. This is achieved by protecting the actual molecular machinery (the enzymes) within the polymersome aqueous lumen away from immunological signalling and proteolytic degradations. We show that such a physical encapsulation enables high flexibility and indeed we show that self-phoresis can be achieved using different combinations of enzymes and substrates, with the only limiting factor being the ability of the substrate to penetrate across the polymersome membrane. We have shown that the combination of glucose oxidase and catalase makes a very efficient chemotactic polymersome in the presence of a glucose gradient. Glucose oxidase and catalase work in tandem to create propulsion, transforming endogenous occurring glucose to endogenous occurring d-glucono-δ-lactone and water, without the formation of potentially harmful compounds such as hydrogen peroxide and gaseous oxygen. Finally, we demonstrate that with very minimal modification, we transform a well established delivery system, the polymersome, into an efficient carrier that enables for the first time the use of chemotaxis to augment biological barrier crossing. This is proved by augmenting the delivery across the blood brain barrier, where we have demonstrated an increase of almost 4-fold in the amount of polymersomes gaining access to the brain parenchyma of rats compared to BBB-targeting, non-chemotactic polymersomes. This is a strong finding that we envision will set a completely new trend in the design of drug delivery systems embracing the new advances being proposed in active colloids.

## Methods

### Materials

Chemicals were used as received unless otherwise indicated. 2-(Methacryloyloxy)ethyl phospho-rylcholine (MPC > 99%) was kindly donated by Biocompatibles, UK. 2-(Diisopropylamino)ethyl methacrylate (DPA) was purchased from Scientific Polymer Products (USA). Copper(I) bromide (CuBr; 99.999%), 2,2-bipyridine (bpy), methanol (anhydrous, 99.8%) and isopropanol were purchased from Sigma Aldrich. The silica used for removal of the ATRP copper catalyst was column chromatography grade silica gel 60 (0.063–0.200 mm) purchased from E. Merck (Darmstadt, Germany). 2-(N-Morpholino)ethyl 2-bromo-2-methylpropanoate (ME-Br) initiator was synthesised according to a previously reported procedure [49]. Poly (ethylene glycol) methyl ether methacrylate P(OEG10MA) was purchased from Sigma Aldrich UK (Dorset, UK). PEO-PBO copolymer was purchased from Advanced Polymer Materials Inc. The polymersomes were labeled using Rhodamine B octadecyl ester perchlorate purchased by Sigma-Aldrich. PBS was made from Oxoid tablets (one tablet per 100 ml of water). Bovine liver Catalase, Glucose Oxidase and glucose have been purchased from Sigma-Aldrich. The gel filtration column for the purification of the polymersomes was made with Sepharose 4B, purchased from Sigma-Aldrich.

### PMPC25-PDPA70 copolymer synthesis

The PMPC-b-PDPA diblock copolymer was prepared by ATRP [49]. In a typical ATRP procedure, a Schlenk flask with a magnetic stir bar and a rubber septum was charged with MPC (1.32 g, 4.46 mmol) and ME-Br initiator (50.0 mg, 0.178 mmol) in ethanol (4 ml) and purged for 30 minutes with N2. Cu(I)Br (25.6 mg, 0.178 mmol) and bpy ligand (55.8 mg, 0.358 mmol) were added as a solid mixture into the reaction flask. The [MPC]: [ME-Br]: [CuBr]: [bpy] relative molar ratios were 25: 1: 1: 2. The reaction was carried out under a nitrogen atmosphere at 20 °C. After 60 minutes, deoxygenated DPA (6.09 g, 28.6 mmol) and methanol (7 ml) mixture were injected into the flask. After 48 h, the reaction solution was diluted by addition of ethanol (about 200 ml) and then passed through a silica column to remove the copper catalyst. The reaction mixture was dialysed against water to remove the organic solvent and then freeze dried. Finally, the copolymer molecular weight was checked by NMR analysis.

### P(OEG10MA)20-PDPA100 copolymer synthesis

The protected maleimide initiator (Mal-Br) was prepared according to a previously published procedure [50] In a typical procedure, either ME-Br or Mal-Br initiators ATRP initiators (0.105 mmol, 1 eq) was mixed with OEG10MA (1 g, 2.11 mmol, 20 eq). When homogeneous, 1 ml water was added, and the solution was purged with nitrogen for 40 minutes. Then, a mixture of CuCl (10.4 mg, 0.105 mmol) and bpy (32.9 mg, 0.210 mmol) was mixed. After 8 minutes, a sample was removed and a nitrogen-purged mixture of DPA (2.2455 g, 0.0105 mol, 100 eq) mixed with 3 ml isopropanol was added to the viscous mixture via cannula. After 18 h, the mixture was diluted with methanol. Then, 2 volumes of dichloromethane were added. The solution was passed through a column of silica using dichloromethane: methanol 2:1 to remove the copper catalyst. The resulting solution was dialysed (MWCO 1,000 Da) against ethanol and water and freeze-dried. The resulting copolymer composition was determined by NMR analysis.

### Copolymer conjugation with cysteine-terminated peptide

The deprotected Mal-P(OEG_10_MA)_20_-PDPA_100_ (105.6 mg, ≃3.4 μmol maleimide) was dispersed in 4.5 ml nitrogen-purged PBS at pH 7.3. The pH was lowered by addition of concentrated HCl (10 μ *l*) to give a uniform solution. The pH was then increased to 7.8 with 5 M NaOH and the resulting opaque dispersion was sonicated for 10 min. 2.3 ml of this solution was transferred to a second flask. Both solutions were then purged with nitrogen for 10 minutes. (This should give an approximate maleimide amount in each flask of 1.7 μmol). To the original solution was then added Cys-Angiopep (5.5 mg, 2.3 μmol thiol) followed by TCEP (2 mg, 7 μmol). The pH in each solution was measured to 7. Both solutions were left for 17 h. Then, both solutions were dialysed against water (MWCO 8,000) to remove any excess peptide, followed by freeze-drying. Successful labelling was confirmed using a HPLC with fluorescence and absorption detection: contains fluorescent tyrosine residues, rendering the polymer-peptide conjugates fluorescent at 303 nm when excited at 274 nm. On the other hand, the non-labelled polymer does not exhibit any fluorescence at these wave-lengths (but can be detected using the absorption detector).

### Polymersome Preparation

Nanometer-sized polymersomes were formed by the film rehydration method [51, 52]. The block copolymers were dissolved in 2:1 v/v chloroform/methanol at 10 mgml^-1^ total copolymer concentration in the organic solvent. Asymmetric polymersomes were obtained by dissolving premixed copolymers at 90% PMPC_25_-PDPA_70_ or P(OEG_10_)MA_20_-PDPA_100_ and 10% PEO_16_-PBO_22_ in molar ratio. Rhodamine B in chloroform solution was added to the above solutions to create a 50 μgml^-1^ fluorophore final concentration. Polymeric films were obtained by drying the copolymer solutions in vacuum oven overnight. In a typical experiment, PBS 0.1 M (pH 7.4) was added to the polymeric films and they were let stir for 30 days at room temperature to obtain the formation of PEO-PBO domains on the PMPC-PDPA polymersomes surface. Topological asymmetry and size distribution have been characterise by TEM and DLS analysis respectively.

### Transmission electron microscopy(TEM)

A phosphotungstenic acid (PTA) solution was used as positive and negative staining agent because of its preferential interaction with the ester groups on the PMPC polymers [53], which are not present in the PEO-PBO copolymer. The PTA staining solution was prepared dissolving 37.5 mg of PTA in boiling distilled water (5 ml). The pH was adjusted to 7.4 by adding a few drops of 5 M NaOH with continuous stirring. The PTA solution was then filtered through a 0.2 μm filter. Then 5 μl of polymersome/PBS dispersion was deposited onto glow-discharged copper grids. After 1 min, the grids were blotted with filter paper and then immersed into the PTA staining solution for 5 s for positive staining, 10 s for negative staining. Then the grids were blotted again and dried under vacuum for 1 min. Grids were imaged using a FEI Tecnai G2 Spirit TEM microscope at 80 kV.

### Dynamic light scattering (DLS)

The sample was crossed by a 120 mW He-Ne laser at 630 nm, at a controlled temperature of 25^®^and the scattered light was measured at an angle of 173°. For the analysis, the sample was diluted with filtered PBS pH 7 at a final concentration of 0.2 mgml^-1^ into a final volume of 500 μ *l* and finally, analysed into a polystyrene cuvette (Malvern, DTS0012). All DLS data were processed using a Dispersion Technology Software (Malvern Instruments).

### Reversed phase high pressure liquid chromatography (RP-HPLC)

RP-HPLC was performed with Dionex Ultimate 3000 instrument equipped with Variable Wavelength Detector (VWD) to analyse the UV absorption of the polymers at 220 nm and the enzymes signal at 280 nm. A gradient of H_2_ O+Tryfluoroacetic acid 0.05% (TFA) (A) and MeOH+TFA 0.05% (B) from 0 min (5%B) to 30 min (100%B) was used to run the samples trough a C18 column (Phenomenex). The peak area was integrated by using Chomeleon version 6.8.

### Enzymes encapsulation

Electroporation was used to allow the entrapment of glucose oxidase, catalase or the combination of the two within the polymersomes. The optimal setting used for the electroporation was 10 pulses at 2500 V [31]. The number of enzymes that can be encapsulated is dictated by the enzyme charge and size. As we demonstrated previously [31], the loading can be modulated changing the electroporation AC voltage intensity, the number of pulses, as well as by adjusting the enzyme surface charges (for example controlling the solution pH). After electroporation, the samples were purified by preparative gel permeation chromatography. Then, the amount of polymer and encapsulated enzymes were quantified by reversed phase high pressure liquid chromatography.

### Encapsulation efficiency calculation

HPLC and DLS data were combined to calculate the number of polymersomes produced in any experiment. The encapsulation efficiency was defined as the number of molecules of enzyme loaded in each polymersomes. The number of polymersomes in a sample can be estimated from the aggregation number (*N*_agg_), defined as:

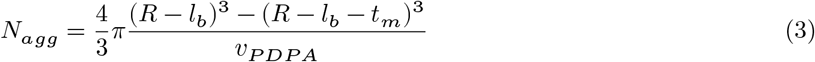

where *R* is the particle radius from the DLS, *l*_b_ is the length of the hydrophilic PMPC brush, *t_m_* is the thickness of the PDPA membrane and *v_PDPA_* is the molecular volume of a single PDPA chain. The number of polymersomes (*N_ps_*) in the sample is defined as

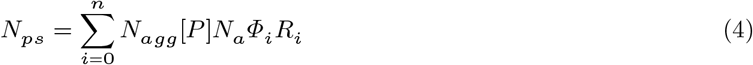

where [*P*] is the moles of copolymer in the sample, *N_a_* is Avogadro’s number and *F_i_.R_i_* is the fraction of sample at a defined radius *R*. Finally, the encapsulation efficiency, *e*, is given by:

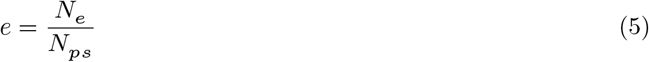

where *N_e_* is the number of enzymes in the sample. The average of encapsulated enzymes per polymersome were 1.9 ± 0.25 for the Catalase and 6 ± 0.45 for the Glucose Oxidase. Results are shown in Supplementary Table 1 and in Supplementary Fig. 1.

### NTA measurements of polymersomes diffusion

Nanoparticle Tracking Analysis (NTA) was performed with a Nanosight^®^ LM14 instrument equipped with a Scientific CMOS camera mounted on an optical microscope to track scattered light by particles illuminated a focused (80 μm) beam generated by a single mode laser diode (405 nm). The polymersomes solution (1 ml) was injected in a concentration of approximately 100 particles/ml in PBS. Samples and controls were injected into the Nanosight^®^ chamber as described in the Supplementary Fig. 2. Two different population of polymersomes (asymmetric and symmetric) were analysed with hydrogen peroxide/glucose, depending on the loaded enzyme. Particles were tracked by the built-in software for 60 seconds at 30 fps. The recorded tracks were analysed using Matlab^®^. Origin of movement for all particles was normalised to Cartesian coordinates (0,0). The mean square displacement (MSD) of all particles was calculated as reported in [5]. Tracks were analysed for 1 s. Particles not tracked for at least 1s were discarded from the analysis. The average number of tracks per sample ranged from 2000 to 10000 traces.

### *In vitro* 3D cell culture blood-brain barrier

For mouse brain endothelial cells (bEnd.3, ATCC^®^ CRL-2299^™^), the medium used was DMEM supplemented with 10% FCS, penicillin and streptomycin, L-glutamine and Fungizone. Astrocyte (ATCC^®^ CRL-2541^™^, C8-D1A Astrocyte Type I clone) medium was antibiotics free DMEM supplemented with 10% FCS and L-glutamine. Pericyte (MSC, Gibco^®^iMouse, C57BL/6) medium used was DMEM F12 media with gluta-MAX-I, supplemented with 10% FCS and 5 gml^-1^ gentamicin. For transwell experiments, both sides of the transwell insert filters (Corning^®^3460 PE filter, diameter: 1.05 cm, pore size: 0.4 m) were pre-coated with 10 gcm^-2^ collagen for 2 hours at room temperature. This was followed by seeding bEnd.3 endothelial cells on the upper surface of the transwell at a density of 20,000–40,000 cells/well, and incubated for 12 hours at 37 °C in 95% air 5% CO2 in order to allow the cells to fully attach. Next, pericytes (10,000–20,000 cells/well) were seeded on the opposite side of the filter insert, and incubated for 12 hours at 37 °C in 95% air 5% CO2. Finally the inserts were moved to a transwell plate, and incubated for 7 days at 37 °C, the medium being changed every two days. Note the medium was supplemented with conditioned medium extracted from astrocyte culture. The endothelial tight juctions were stained either with anti ZO^-1^ and Claudin-5, while pericytes are shown using anti-CD140. For confocal imaging, the BBB models is fixed and imaged using a stack of 100 images with an optical slice of 0.4pm. The concetration of polymersomes on the upper (apical) and lower (basolateral) compartments are measured by HPLC using a fluorecence detectors collecting samples at different time points. For the early time point and live cell kinetics, brian endothelial cells were treated with CellMASK^®^ for 30mins, washed 3 times with PBS and an immersed in imaging media (FluoroBrite™ DMEM) supplemented with 10% FCS and 5 gml^-1^ gentamicin. Polymersomes were subsequently added at a concentration of 1 mg/ml into the apical (upper) transwell compartment after Trans-Epithelial Electric Resistance (TEER) measurements were taken with an EVOM2 Epithelial Voltohmmeter. Cells were incubated for 1–2 hours at 37 °C in 95% air 5% CO2 and were imaged on Leica SP8 confocal laser-scanning microscope with 40x water immersion lens and 63x oil immersion lens. Rhodamine-labelled polymersomes, an excitation energy 561 nm was used and fluorescence emission was measured at 575–600 nm. Membrane staining was performed using CellMASK^®^. Image data was acquired and processed using Image J software. We repeated this experiment three times and measured a total of 35 crossing events

### Brain *in situ* perfusion

All animal experiments were performed in accordance with the Animals (Scientific Procedures) Act 1986 (U.K.) Male adult Wistar rats were anaesthetised with 100 mgkg^-1^ ketamine and 1 mgml^-1^ medetomidine via intraperitoneal injection. The right and left external carotid arteries were isolated from the carotid sheaths and cannulated according to a previously established procedure [54]. The perfusion fluid was modified Ringer’s solution (6.896 g l^-1^ NaCl, 0.350 g l^-1^ KCl, 0.368 gl^-1^ CaCl2, 0.296 g l^-1^ MgSO4, 2.1 g l^-1^ NaHCO3, 0.163 g l^-1^ KH_2_O_4_, 2.383 g l^-1^ HEPES, additionally 0.5005 g l^-1^ glucose (5.5 mM) and 11.1 g l^-1^ BSA). The perfusion fluid was bubbled with 5% CO_2_ and heated to 37 °C for 20 minutes prior to perfusion. For the injection of polymersomes, 20% (mol) Cy3-labelled polymersomes in PBS with our without protein encapsulated were diluted to 1 mg ml^-1^ in Krebs buffer (pH 7.4, 188 mM NaCl, 4.7 mM KCl, 2.5 mM CaCl_2_, 1.2 mM MgSO_4_, 1.2 mM KH_2_PO_4_, 25 mM NaHCO_3_, 10 mM D-glucose, 3 g l^-1^ BSA). The polymersome solution was supplied via syringe pump at 0.16 ml min^-1^, with a total perfusion rate of 1.5 ml min^-1^ and a total perfusion time of 10 min. At the end of the perfusion time, the syringe pump was stopped and the arteries were flushed for 60 s with modified Ringer’s perfusate in order to remove unbound polymersomes. After 60 s, cerebrospinal fluid was extracted via cisternal puncture followed by decapitation and removal of the brain.

### Quantification of polymersome distribution in the rat brain

After decapitation, brains were removed and washed in ice cold 9 gl^-1^ NaCl, followed immediately by homogenisation on ice to initiate the capillary depletion method [54]. Briefly, the cerebellum was removed and the cerebrum was weighed, adding 2x brain weight in PBS followed by 3x dilution in 30% (w/v) dextran (average **MW** 64–74 kDa). Centrifugation of homogenates at 7400g for 20 minutes in 4°C resulted in several fractions that were carefully separated: capillary depleted (CD) fraction (i.e. parenchyma), dextran, and the capillary enriched fraction (pellet). The capillary enriched pellet was re-suspended in PBS, and 100 μ L samples were added to a black 96-wellplate and read in a fluorimeter at an excitation wavelength of 540 nm and emission at 565 nm. All sample fluorescence readings were normalised to readings obtained from sham perfused rats (n=6) for each sample type, i.e. CD, dextran or capillaries. Positive controls were polymersomes in perfusate harvested from the cannula at the injection point. Normalised fluorescence readings were converted to polymersome (Cy3) amount was converted into percentage injected dose %id of the positive control value for that experiment, where %id = [normalised sample value (mg) / mean positive control value (mg)]* 100. This was further converted into fluorescence per whole brain. All statistical analysis was one-way ANOVA, p <0.05. All animal studies were carried out according to the ARRIVE guidelines under licence from the UK Home Office, (Scientific Procedures Act 1986) and approved by the King’s College London Ethical review committee.

## Acknowledgements

We would like to thank Prof Ramin Golestanian from Oxford University for the valid and critical discussions at the early stages of this work, Profs Anthony Ryan and Steve Armes from Sheffield University for supplying some of the copolymers we used for the initial validation work. We thank Dr Tung Chun Lee from the UCL Institute for Materials Discovery and Prof Giovanni Volpe from Bilkent University for reading our manuscript and providing us with valid comments. We acknowledge the ERC for the MEViC ERC-STG project that supported most of the experimental work and the salary of A.J. and L.R-P. as well as part of J.G. and G.B. salary, and the EndoNaut ERC-PoC project for supporting D.C. salary. J.G. thanks the Deutsche Forschungsgemeinschaft (DFG) for supporting his postdoctoral fellowship, C.C., S.N. and G.F. are thankful to the UCL MAPS faculty for sponsoring their studentship. G.V. acknowledges partial financial support by the COST Action MP1305.

## Author contributions

G.B., D.C., A.J., and C.C. designed the *in vitro* experiments and performed the data analysis, G.B, S.N. and J.P. designed the animal experiments and performed the data analysis, A.J. designed and performed the computational fluid dynamics simulations, G.F. designed and performed the agent-based simulations, G.V. designed the numeric simulations. A.J. performed most of the numeric simulations. G.B, G.V., A. J. and G.F. analysed the simulation data. D.C. performed all the preliminary *in vitro* experiments, C.C. performed all the final *in vitro* experiments including DLS, TEM and encapsulation experiments. X. T. optimised the *in vitro* BBB model and together with S.N. performed the BBB crossing experiments. S.N., J.A and J.P. performed all the animal experiments. J.G. synthesised all the PMPC-PDPA and P(OEGMA)-PDPA copolymers used in the work. L.R-P. analysed the initial samples by transmission electron microscopy and designed the protocols for selective staining. GB, D.C., A.J., G.V., L.R-P., G.F., S.N. and C.C. wrote the manuscript

